# Intra-host symbiont diversity and extended symbiont maintenance in photosymbiotic Acantharea (clade F)

**DOI:** 10.1101/299495

**Authors:** Margaret Mars Brisbin, Lisa Y. Mesrop, Mary M. Grossmann, Satoshi Mitarai

## Abstract

Photosymbiotic protists contribute to surface primary production in low-nutrient, open-ocean ecosystems and constitute model systems for studying plastid acquisition via endosymbiosis. Little is known, however, about host-symbiont dynamics in these important relationships, and whether these symbioses constitute mutualisms is debated. In this study, we applied single-cell sequencing methods and advanced fluorescent microscopy to investigate host-symbiont dynamics in clade F acantharians, a major group of photosymbiotic protists in oligotrophic subtropical gyres. We amplified the 18S rRNA gene from single acantharian hosts and environmental samples to assess intra-host symbiont diversity and to determine whether intra-host symbiont community composition directly reflects the available symbiont community in the surrounding environment. Our results demonstrate that clade F acantharians simultaneously host multiple species from the haptophyte genera *Phaeocystis* and *Chrysochromulina*. The intra-host symbiont community composition was distinct from the external free-living symbiont community, suggesting that these acantharians maintain symbionts for extended periods of time. After selective staining of digestive organelles, fluorescent confocal microscopy showed that symbionts were not being systematically digested, which is consistent with extended symbiont maintenance within hosts. Extended maintenance within hosts may benefit symbionts through protection from grazing or viral lysis, and therefore dispersal, provided that symbionts retain reproductive capacity.

## 1 Introduction

Photosymbiosis, a nutritional symbiosis where a heterotroph hosts photosynthetic endosymbionts, substantially increases surface primary production in oligotrophic marine ecosystems (Not et al. 2016). Photosymbiosis is also believed to have led to evolution of eukaryotic oxygenic photosynthesis and eventual emergence of diverse photosynthetic eukaryotes, with many evolving from eukaryote-eukaryote secondary and tertiary endosymbioses (Keeling 2004). Eukaryote-eukaryote photosymbioses continue to be extremely common among marine protists and contribute significantly to the productivity of oligotrophic open-ocean ecosystems (Decelle et al. 2015). Nonetheless, little is known about host-symbiont dynamics, such as host-symbiont specificity or host mechanisms for symbiont recognition, uptake, and maintenance. Whether photosymbioses are mutualistic continues to foster debate, because the exact nature of these relationships may explain how they evolve and persist (Keeling & McCutcheon 2017).

The Acantharea belong to the Rhizaria, a large group of amoeboid protists that includes many photosymbiotic lineages (Burki & Keeling 2014). Photosymbiotic acantharians are often the most abundant photosymbiotic Rhizaria in oligotrophic surface waters (Michaels et al. 1995), where they contribute significantly to primary production (Caron et al. 1995). The majority of acantharian species (clades E and F) host algal symbionts in the Haptophyte genus *Phaeocystis* (Decelle et al. 2012). *Phaeocystis* is a globally distributed genus with species that present multiple phenotypes— solitary, flagellated, and colonial—and sometimes form harmful algal blooms (Schoemann et al. 2005). Despite the ecological significance of both partners, this symbiosis remains largely unstudied. There is some evidence, however, suggesting that this relationship is not a case of mutualism and symbionts are instead exploited (Decelle et al. 2012).

Traditionally, photosymbioses are considered mutualisms under the assumption that hosts provide nitrogen to symbionts and symbionts provide organic carbon to hosts in return (Garcia & Gerardo 2014). Whether these relationships are truly mutualistic or are instead cases of symbiont exploitation has been increasingly questioned in recent years and there is mounting evidence that exploitation is the rule rather than the exception (Keeling & McCutcheon 2017). In high-light and low-prey conditions, as found in oligotrophic surface waters, protistan photosymbiotic hosts benefit from an increased growth-rate, but symbiont growth-rate is suppressed and photosynthetic efficiency is decreased compared to free-living symbionts. These results indicate that algal symbionts may actually experience restricted nitrogen availability *in hospite* and do not benefit from symbiosis (Lowe et al. 2016). Estimated free-living populations of *Phaeocystis* in oligotrophic conditions (Moon-van der Staay et al. 2000) are much larger than possible symbiotic populations estimated from acantharian abundance and symbiont load (Michaels 1991). Symbiont growth rate may therefore be decreased within acantharian hosts, potentially indicating that Acantharea-*Phaeocysti*s photosymbioses are also exploitative rather than mutualistic (Decelle 2013).

Increased growth rate is not the only means by which a symbiont may benefit from a host: decreased predation, viral attack, or competition *in hospite* may allow symbionts to benefit from enhanced dispersal and future reproduction, assuming reproductively viable symbionts are released from hosts (Douglas 2010; Garcia & Gerardo 2014). Reproducing symbionts are known to be released from some photosymbiotic protistan hosts: *Chlorella* escapes from *Paramecium* hosts and establishes free-living populations when low-light inhibits host benefit (Lowe et al. 2016) and dinoflagellate symbionts of colonial radiolarians establish free-living populations when isolated from hosts (Probert et al. 2014). Some photosymbiotic forams, however, digest all of their symbionts prior to gametogenesis (Takagi et al. 2016). It is currently unknown whether symbiotic *Phaeocystis* retains reproductive capacity, but symbiotic cells have not yet been cultured from hosts (Decelle et al. 2012). It is possible that phenotypic changes observed in symbiotic *Phaeocystis*—additional plastids and an enlarged central vacuole (Febvre et al. 1979)—are evidence that symbionts are incapable of cell division, which would make the relationship an ecological and evolutionary dead-end for *Phaeocystis* and preclude the possibility for mutualistm (Decelle et al. 2012).

The number of symbionts observed in individual acantharians increases with host size (Michaels 1991), thus requiring that symbionts reproduce *in hospite*, that acantharians recruit new symbionts, or possibly both. If acantharians recruit one or a few symbionts early in development and then support a reproducing symbiont community, the intra-host symbiont community would exhibit low diversity and may be divergent from the free-living environmental community. If symbionts divide within hosts, hosts must exert an alternative form of population control, potentially by shedding (mutualism) (Boettcher et al. 1996; Fishman et al. 2008) or digesting (exploitation) excess symbionts (Titlyanov et al. 1996). Conversely, if acantharians recruit new symbionts, the intra-host symbiont community is likely to be more diverse and representative of the available free-living symbiont community in the surrounding waters. Low-diversity intra-host symbiont communities would therefore suggest that symbionts maintain reproductive capacity and allows for possible symbiont benefit, whereas high diversity communities may be interpreted as further evidence against mutualism. However, neither intra-host symbiont diversity, nor the relationship between symbiont identity and environmental availability of potential symbionts have been investigated in clade E or F acantharians.

In this study, we applied single-cell Next Generation Sequencing (NGS) to illuminate intra-host symbiont diversity in individual clade F acantharians collected from 7 sampling sites along the Ryukyu Archipelago, spanning more than 1,000 km in the East China Sea (ECS), and near Catalina Island (California, USA). We further applied NGS to evaluate the environmental availability of symbionts to acantharian hosts sampled from the ECS. We compared molecular symbiont diversity with intra-host symbiont population size by enumerating symbionts with fluorescent confocal microscopy in a subset of acantharians prior to nucleic acid extraction. Additional acantharians were collected and imaged after selectively staining lysosomes and phagolysosomes in order to observe their proximity to symbionts and to determine if symbionts are systematically digested. This study provides new evidence to the mutualism-exploitation debate relative to Acantharea-*Phaeocystis* symbioses by investigating intra-host symbiont diversity and by assessing host-symbiont specificity in the context of environmental symbiont availability.

## 2 Materials & Methods

### 2.1 Individual acantharian sampling

Single acantharians were collected from coastal water near Catalina Island (California, U.S.A.), Okinawa Island (Okinawa, Japan), and from cruise stations visited during the Japan Agency for Marine-Earth Science and Technology (JAMSTEC) MR17-03C cruise to the ECS aboard the R/V *Mirai* in May and June 2017 (Figure 1, Supplementary Table S1). Okinawa Island and Catalina Island plankton samples were collected by pulling a Rigo Simple 20 cm diameter, 100-μm-mesh plankton net or a SEA-GEAR 12” diameter, 163-μm-mesh plankton net, respectively, along the sea surface approximately 5 m behind a small craft at its lowest speed. Aboard the R/V *Mirai*, plankton samples were collected by passing unfiltered seawater pumped from the sea surface through a 100-μm-mesh, hand-held plankton net (Rigo). Plankton samples were observed under a dissecting microscope and individual acantharians were transferred by glass micropipette to clean petri-dishes. Acantharians were rinsed with 0.2-μm-filtered seawater several times until all visible contaminants were removed and were then incubated for 0.5–2 hr to allow additional self-cleaning. Acantharians collected aboard the R/V *Mirai* and those from near Okinawa Island were imaged with inverted light microscopy (Zeiss Primovert, Olympus CKX53, Supplementary Figures S1 and S2). Several acantharians collected near Okinawa Island were also imaged with laser confocal microscopy (described below). Each acantharian was transferred to a maximum recovery PCR tube (Axygen) and successful transfer was confirmed by microscopy before adding 30 μL of RLT-plus cell-lysis buffer to each tube (Qiagen). Immediately following buffer addition, samples were flash-frozen with liquid nitrogen and stored at -80°C until later processing in the lab.

**Figure 1.**
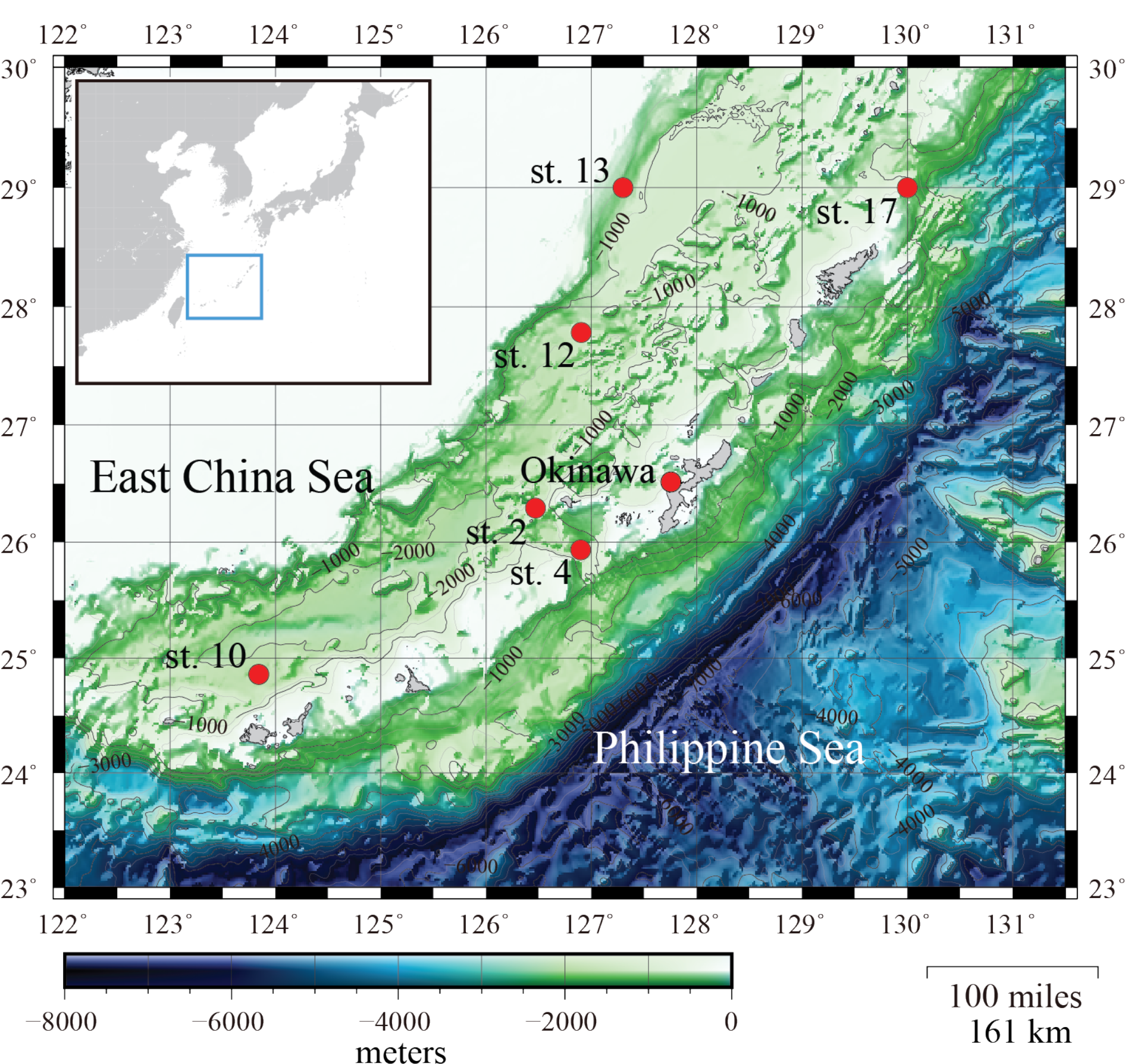
Acantharea and seawater samples were collected along the Ryukyu Archipelago, spanning 1,000 km in the East China Sea (ECS). Stations 2, 4, 10, 12, 13, and 17 were sampled during the Japan Agency for Marine-Earth Science and Technology (JAMSTEC) MR17-03C cruise to the ECS aboard the R/V *Mirai* in May and June 2017. Samples were collected from the Okinawa Island (Okinawa, Japan) sampling site in April, May, and December 2017. Additional samples were collected near the University of Southern California’s Wrigley Institute for Environmental Studies on Catalina Island (California, U.S.A., GPS: 33°26′59.12″N 118°29′17.17″W) in May 2017. Bathymetry map is courtesy of JAMSTEC.

### 2.2 Environmental sampling

Seawater samples were collected at each ECS cruise station visited by the JAMSTEC MR17-03C cruise where acantharians were also isolated. Two replicates of 4.5 L of seawater were collected from the sea surface by bucket and sequentially filtered under a gentle vacuum through 10.0 μm and 0.2 μm pore-size Polytetrafluoroethylene (PTFE) filters (Millipore) to size-fractionate plankton and separate free-living *Phaeocystis* (< 10 μm) from acantharian hosts (> 10 μm). Filters were flash-frozen in liquid nitrogen onboard and stored at -80°C until processing onshore.

### 2.3 RNA extraction from individual acantharian hosts

RNA extractions from single acantharians (n = 42) were accomplished by modifying methods of Trombetta et al. (2015). Samples were thawed over ice, vortexed twice (10 s, speed 7, Vortex-Genie 2), and then incubated at room temperature for 5 min to fully lyse cells. Agencourt RNAClean XP magnetic beads (Beckman Coulter) were added to each sample at a 2.2 : 1 v:v ratio and fully mixed by pipette prior to a 30-min incubation in order to bind all RNA to the magnetic beads. After two 80% ethanol washes, RNA was eluted from the beads in 11 μL of a custom elution buffer (10.72 μL nuclease-free water, 0.28 μL RNAase inhibitor (Clonetech)) and 10.5 μL of eluted RNA was further processed following the single-cell protocol for the SMART-seq v4 Ultra Low Input Kit (Clonetech) with 18 cycles in the primary PCR. The resulting cDNA from each sample was quality checked with the Bioanalyzer High Sensitivity DNA Assay (Agilent) and quantified with the Qubit dsDNA High Sensitivity Assay (Qubit 3.0, ThermoFisher).

### 2.4 DNA extraction from environmental samples

Environmental DNA was extracted from the 0.2-μm pore-size PTFE filters (n = 12, 2 replicates from 6 stations) following manufacturer protocols for the Qiagen AllPrep DNA/RNA Mini Kit with limited modifications. Half of each PTFE filter was submerged in RLT-plus cell-lysis buffer (Qiagen) with garnet beads in a 2-mL tube (MoBio/Qiagen). Samples were heated for 10 min at 65°C and then vortexed at maximum speed (Vortex-Genie 2) for 5 min with the MoBio/Qiagen vortex adapter to fully lyse cells. After cell lysis, DNA extraction proceeded without further modifications. Extracted DNA was quantified with the Qubit dsDNA High Sensitivity Assay on a Qubit 3.0 instrument (ThermoFisher).

### 2.5 Library preparation and sequencing

Library preparation for acantharian cDNA samples and environmental DNA samples followed procedures described in the Illumina 16S Metagenomic Sequencing Library Preparation manual, modified only to include universal eukaryotic primers for the v4 region of the eukaryotic 18S rRNA gene (Stoeck et al. 2010) and amplicon PCR conditions most appropriate for these primers. The forward primer, TAReuk454FWD1 (CCAGCASCYGCGGTAATTCC, Stoeck et al. 2010), was used unmodified. The reverse primer, TAReuk454REV3 (ACTTTCGTTCTTGATYRA, Stoeck et al., 2010) was reported not to amplify the *Phaeocystis* 18S gene in *in silico* PCRs (Tanabe et al. 2016). Further investigation revealed a mismatch between the final 3’ adenine in the primer sequence and the *Phaeocystis* 18S gene sequence. Although we found that the original primers do amplify the v4 region of the *Phaeocystis* 18S gene in *de facto* PCRs with DNA extracted from *Phaeocystis* cultures, the mismatch could create bias against *Phaeocystis* sequences in more diverse samples. We therefore eliminated the 3’ “A” in the TAReuk454REV3 sequence and used a new reverse primer TAReuk454REV3.1 (ACTTTCGTTCTTGATYR). The optimum annealing temperature for the Illumina-adapted primers was determined by performing temperature gradient PCRs (53–65°C, 0.5°C steps) and the annealing step in the amplicon PCR was set at 58°C thereafter. Following the second, indexing PCR and final product purification, amplicon libraries were quantified with the Qubit dsDNA High Sensitivity Assay (Qubit 3.0, ThermoFisher) and the amplicon size was determined with the Bioanalyzer High Sensitivity DNA Assay (Agilent). Amplicon libraries were then submitted to the Okinawa Institute of Science and Technology (OIST) sequencing center for 300×300-bp paired-end sequencing on the Illumina MiSeq sequencing platform with v3 chemistry.

### 2.6 Amplicon sequence analysis and annotation

Demultiplexed paired-end sequences provided by the OIST sequencing center were imported to Qiime2 (v2017.11) software (Caporaso et al. 2010, qiime2.org). The Divisive Amplicon Denoising Algorithm (DADA) was implemented with the DADA2 plug-in for Qiime2 for quality filtering, chimera removal, and feature table construction (Callahan et al. 2015). DADA2 models the amplicon sequencing error in order to infer sample composition more accurately than traditional Operational Taxonomic Unit (OTU) picking methods (Callahan et al. 2015). Taxonomy was assigned to the feature table in Qiime2 by importing SILVA 18S representative sequences and consensus taxonomy (release 128, Quast et al. 2013). The feature table and associated taxonomy were then extracted from Qiime2 for further analysis in the R statistical environment (R Core Team 2013). The feature table was split into a table of acantharian samples and a table of environmental samples and both were converted to phyloseq objects for preprocessing and analysis (McMurdie & Holmes 2013). Prevalence filtering was applied to the acantharian feature table to remove low-prevalence (< 5%) SVs and decrease the chance of data artifacts affecting the analysis (Callahan et al. 2016). Prevalence filtering effectively eliminated sequences from known Rhizaria prey (i.e. metazoans and diatoms (Swanberg & Caron 1991)) and parasites (i.e. alveolates (Bråte et al. 2012)). The phyloseq object was further filtered to eliminate SVs originating from hosts by retaining SVs classified as belonging to the phylum Haptophyta, which includes *Phaeocystis*, leaving 21 symbiotic SVs. The environmental feature table was then filtered to include only the 21 symbiotic SVs found in acantharian samples.

The 21 symbiotic SVs were further classified by building a phylogenetic tree. An initial BLAST query (BLASTn 2.8.0+, 03/23/2018, Camacho et al. 2009) indicated that the symbiont SVs belong to the Haptophyte genera *Phaeocystis* and *Chrysochromulina*. Likewise, a SILVA SSU sequence search (03/23/2018, Quast et al. 2013) classified 18 of the sequences as class Prymnesiophycae, order Phaeocystales or Prymnesiales, and genus *Phaeocystis* or *Chrysochromulina*, when classified to genus level. The remaining 3 symbiotic SVs were designated “unclassified” in the SILVA sequence search. Reference 18S rRNA gene sequences were downloaded from GenBank (Benson et al. 2012) for the 5 Haptophyte orders (Pavlovales, Phaeocystales, Prymnesiales, Isochrysidales, and Coccolithales (Edvardsen & Medlin 2007)) and included in a Multiple Sequence Comparison by Log Expectation (MUSCLE) v3.8.31 (Edgar 2004) alignment along with the 21 symbiotic SVs. A Bayesian phylogenetic tree was then built from the alignment using MrBayes v3.2.7 (Ronquist & Huelsenbeck 2003) (Figure 3). The phylogenetic relationships between the Haptophyte orders in the resulting tree match published phylogenies for Haptophyta (Edvardsen & Medlin 2007; Sáez et al. 2004). Similarly, the phylogenetic relationships between cultured clades of *Phaeocystis* matched published phylogenetic trees (Andersen et al. 2015), and the placement of the uncultured clade, Phaeo2, is similar to published single-gene phylogenetic reconstruction with the 18S rRNA gene (Decelle et al. 2012). A phylogenetic tree was built for the 5 dominant acantharian host SVs following the same methods (Figure 4) and the reconstruction matched published phylogenetic relationships for clades E and F Acantharea (Decelle et al. 2012b).

**Figure 3.**
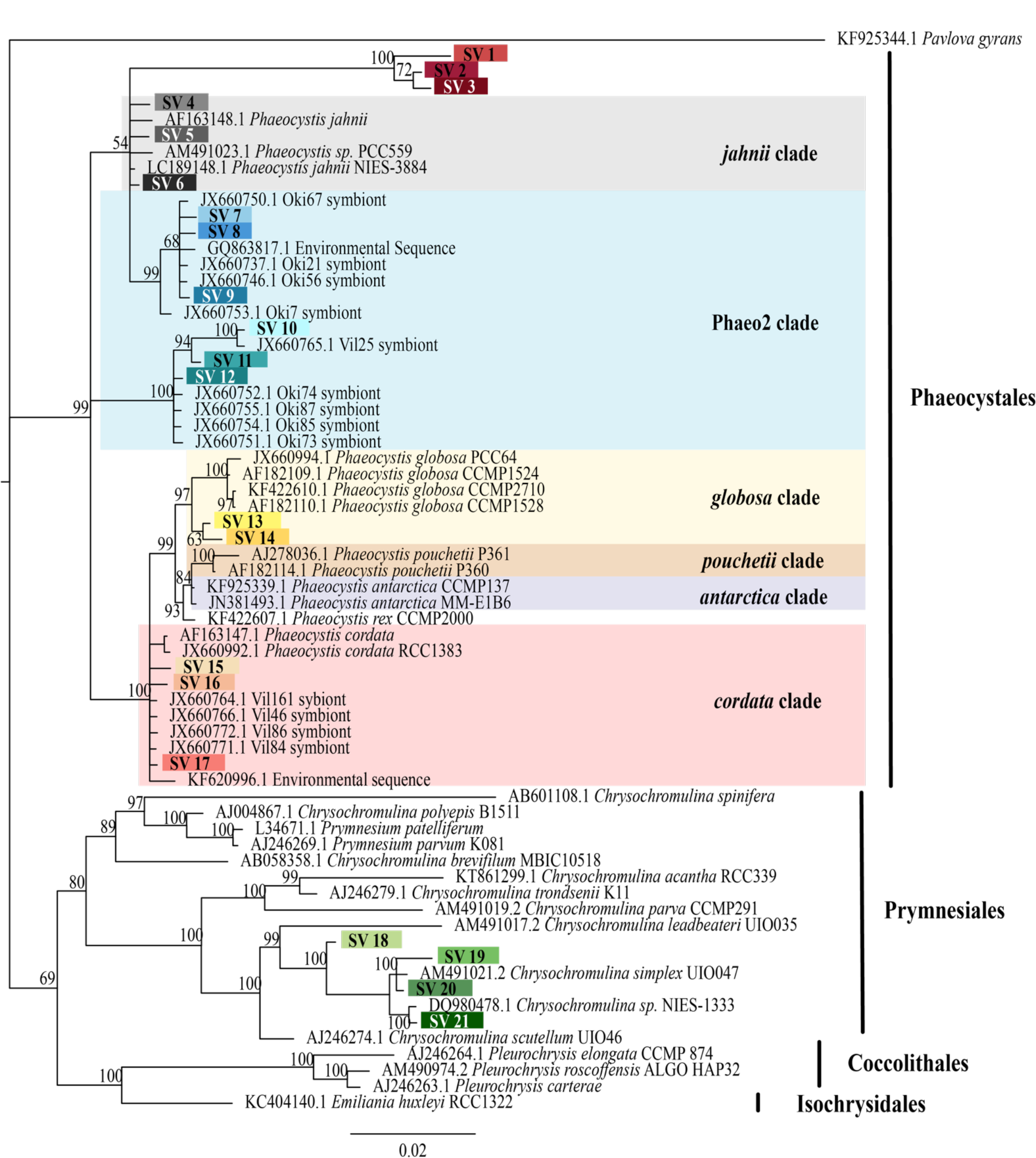
Symbiotic SVs from individual acantharians belong to Haptophyte genera *Phaeocystis* and *Chrysochromulina*. Phylogenetic tree showing the taxonomy of the 5 Haptophyte orders including the 21 symbiotic SVs identified in this study. The symbiotic SVs are highlighted to match the key in Figures 2 and 7. *Phaeocystis* clades are color coded and include the Phaeo2 clade, which is an uncultured clade identified from symbiotic sequences from acantharians collected near Okinawa (Decelle et al. 2012). The tree was built with MrBayes v3.2.7 using a MUSCLE v3.8.31 alignment of the 21 SVs and reference sequences downloaded from GenBank. *Pavlova gyrans* was designated as the outgroup. Values associated with nodes are posterior probabilities as a percent after 100,000 generations (average SD of split frequencies = 0.019). The scale bar indicates 0.02 changes expected per site.

**Figure 4.**
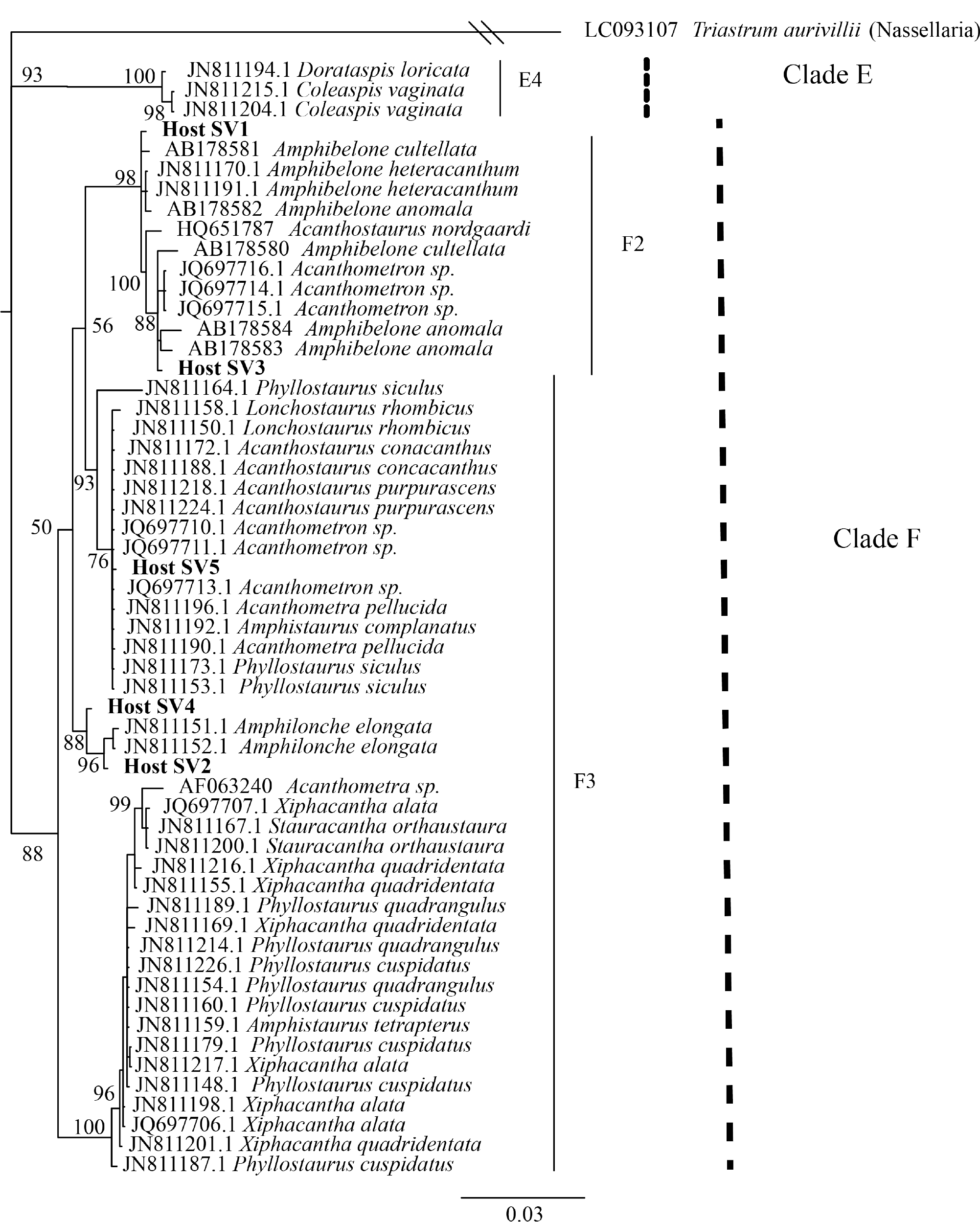
Host SVs belong to 3 genera of clade F Acantharea. Phylogenetic tree including acantharian host SVs and acantharian reference sequences. The tree was built with MrBayes v3.2.7 using a MUSCLE v3.8.31 alignment of the 5 host SVs and reference sequences downloaded from GenBank. The nasellarian radiolarian *Triastrum aurivillii* served as the outgroup. Values associated with nodes are posterior probabilities as a percent after 40,000 generations (average SD of split frequencies = 0.01). The scale barindicates 0.03 changes expected per site.

### 2.7 Statistical analyses

Bray-Curtis distances between symbiont community compositions in acantharian and environmental samples were computed from relative abundances of symbiotic SVs in the filtered feature tables with the R package phyloseq (McMurdie & Holmes 2013). Bray-Curtis distances were used to perform Principal Coordinate Analyses (PCoA) within the phyloseq package, and PCoA plots were rendered with the R package ggplot2 (Wickham 2009). Permutational Multivariate Analyses of Variance (PERMANOVA) with 999 permutations were performed with the adonis function in the R package vegan (Oksanen et al. 2013) to determine whether clustering observed in the ordination plots was statistically significant and to discern which covariates were deterministic of symbiont community composition. PERMANOVA were also performed with the beta-group-significance function in the Qiime2 diversity plugin, which allows for pairwise comparisons. Differences were considered statistically significant when the p-value was ≤ 0.05.

### 2.8 Fluorescent confocal microscopy

In order to observe and enumerate symbionts within hosts for which symbiont communities were also evaluated, acantharians collected near Okinawa in April (Oki.3A, Oki.4A) and May (Oki.3, Oki.7, Oki.10, Oki.11) were imaged without staining using an inverted laser scanning confocal microscope (Zeiss LSM 780) prior to RNA extraction. A z-stack of chlorophyll autofluorescence (em670 nm) and halogen light images was assembled for each host to compare the number of visually observable symbionts to the number of symbiotic SVs identified. In order to evaluate possible host digestion of symbionts, additional acantharian samples were collected in December 2017 (n = 3) and stained with LysoTracker Green DND-26 (ThermoFisher), a fluorescent dye that selectively stains acidic compartments (i.e. digestive organelles) within cells, including lysosomes and phagolysosomes. The LysoTracker dye was diluted from the 1-mM stock solution to a 100-nM working solution in 0.2-μm- filtered seawater and each sample was incubated in 100 μL working solution in the dark for 2 hr before imaging. Z-stacks for these samples were assembled by imaging with 3 channels: red for autofluorescence from symbiont chlorophyll (em670 nm), green for LysoTracker-stained host lysosomes and phagolysosomes (em511 nm), and grey for halogen light imaging. FIJI Image-J software (Schindelin et al. 2012) was used to adjust image brightness, merge color channels, and create 3-D projections for all imaged host cells.

## 3 Results

### 3.1 Intra-host symbiont diversity in individual acantharians

To determine if and to what extent intracellular symbiont diversity exists in acantharians, we performed single-cell RNA extractions and sequenced 18S rRNA gene amplicons from 42 individual acantharians. A total of 6,154,808 sequences were generated from acantharian samples, with 28,828– 241,809 sequences per sample. After quality filtering and feature table construction, 3,294,093 total sequences remained (5,579–136,339 per sample). Within each acantharian sample, 1–17% of sequences derived from symbionts. We identified 21 symbiotic SVs in the acantharian samples, and each acantharian host contained 4–12 unique symbiotic SVs (mean = 8, standard deviation = 2) (Figure 2). Our phylogenetic analysis determined that symbiotic SVs belonged to four *Phaeocystis* clades: the *globosa* clade, the *cordata* clade, the *jahnii* clade, which has not previously been reported as a symbiont, and the uncultured Phaeo2 clade, which was discovered by Decelle et al. (2012) as an acantharian symbiont near Okinawa (Figure 3). Additionally, four symbiotic SVs belonged to the genus *Chrysochromulina*, which had not previously been identified as a symbiont in clade F acantharians. The majority of symbiotic SVs in acantharians collected from the ECS belonged to the *Phaeocystis* clades *cordata*, *jahnii* and Phaeo2, and only 3 of these samples contained SVs belonging to *Chrysochromulina*. The opposite pattern was observed in samples collected near Catalina Island: the majority of symbiotic SVs in these samples were *Chrysochromulina spp*. However, all three samples from Catalina Island hosted symbionts from the Phaeo2 *Phaeocystis* clade, which had previously only been found in hosts collected near Okinawa Island (Decelle et al. 2012). These results demonstrate that acantharians simultaneously host multiple symbiont species and that *Phaeocystis*-hosting acantharians can also host *Chrysochromulina spp*.

**Figure 2.**
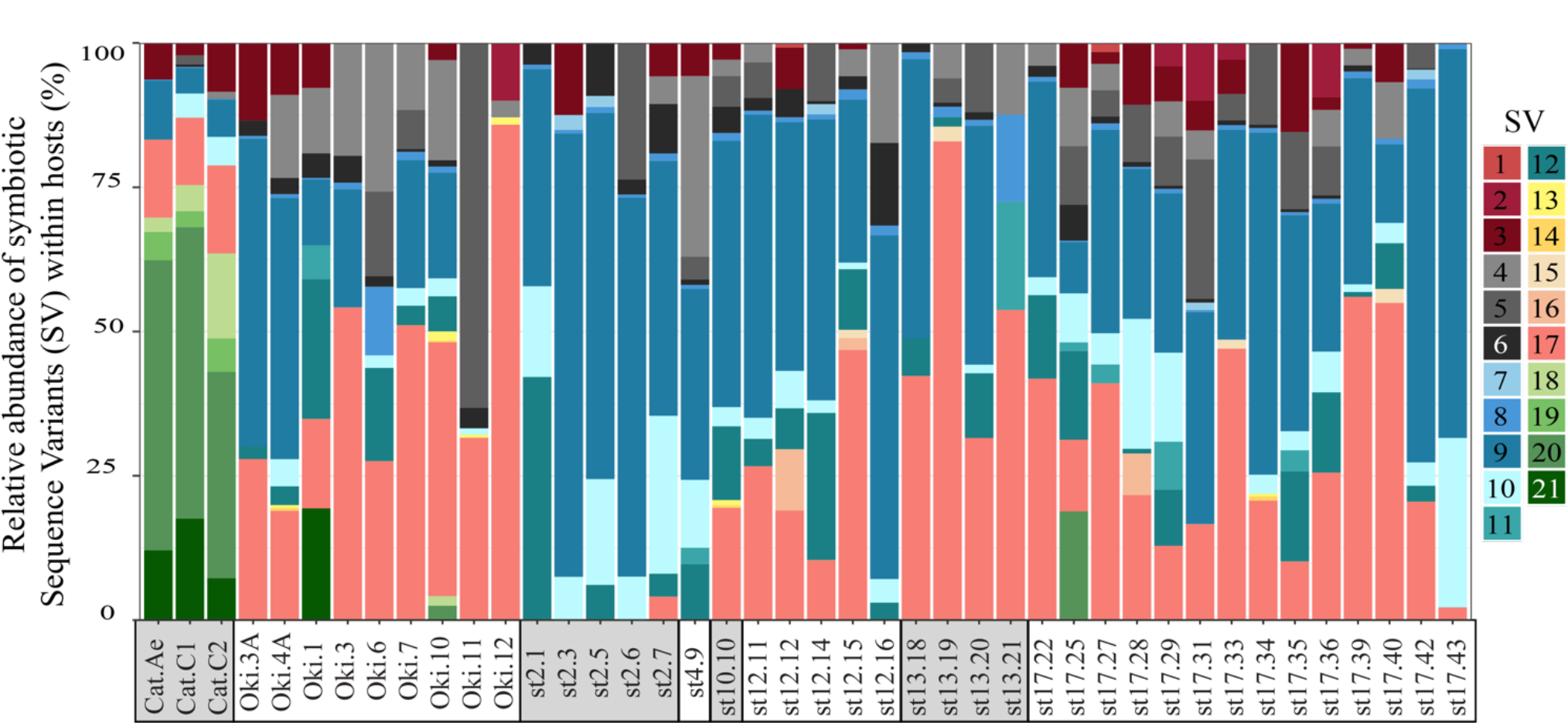
Individual acantharian samples contain multiple Haptophyte Sequence Variants (SVs). The stacked bar graph represents the relative abundance of 21 Haptophyte SVs that were observed in individual acantharian samples. Each bar represents a single acantharian host and is labeled by collection location (Cat: Catalina Island, Oki: Okinawa Island, st#: ECS cruise station number) and sample ID. Individual acantharians contained 4-12 symbiotic SVs (mean = 8, SD = 2). The figure was created with R package ggplot2.

We also investigated the effect of host type and collection location on symbiont community composition within individual acantharians. Acantharian samples were each dominated by 1 of 5 unique Rhizarian SVs, all of which belonged to Acantharea clade F and represented 3 genera: *Amphibelone* (Host SVs 1 and 3), *Amphilonche* (Host SVs 2 and 4), and *Acanthometra* (Host SV5) (Figure 4). Host SV did not appear to determine the symbiont community in a Principal Coordinate Analysis (PCoA) plot based on Bray-Curtis distances between acantharian symbiont community compositions, but collection location appeared to play a role and the Catalina Island samples formed a distinct cluster in the PCoA plot (Figure 5). Statistical testing showed that collection location had a significant effect on symbiont community (p = 0.001, R^2^ = 0.47) and in pairwise comparisons, only two comparisons—st. 17 to st. 12 and st. 13 to Okinawa—were not significantly different (Supplementary Table S2). Host SV also significantly affected symbiont community (p = 0.009, R^2^ = 0.19), but pairwise comparisons showed that the significance was driven solely by the symbiont community associated with host SV 5, which was only found near Catalina Island, indicating that host effect was confounded by location (Supplementary Table S3).

**Figure 5.**
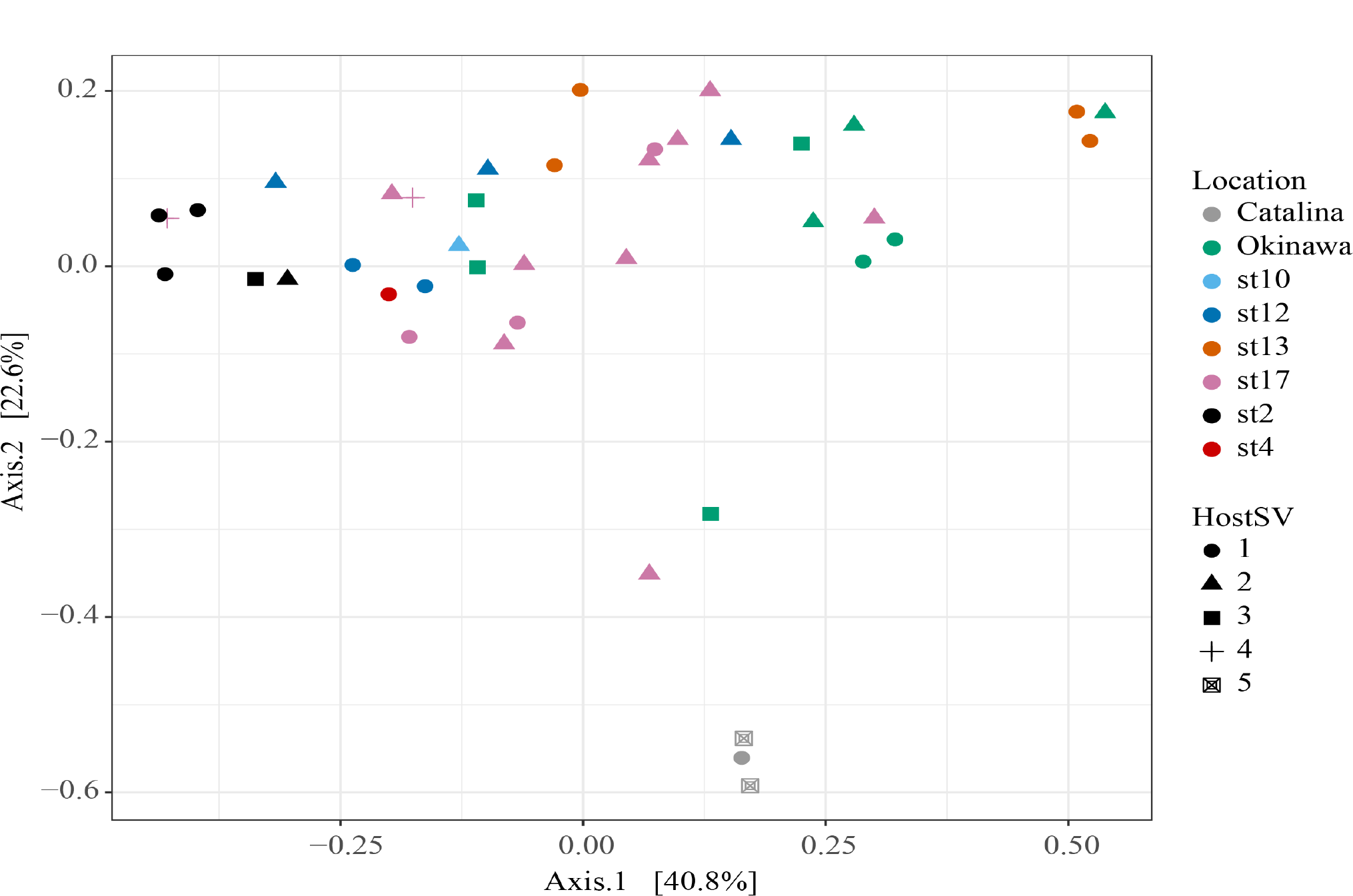
Host collection location has larger effect than host type on the intra-host symbiont community. PCoA plot from Bray-Curtis distances between acantharian symbiont community compositions (n = 42). Point shape corresponds to host sequence variant (SV) and point color corresponds to the collection location of the host acantharians. The symbiont communities associated with acantharians collected near Catalina Island and from ECS station 2 form clusters while communities from the other locations do not cluster separately. A PERMANOVA by location (excluding st. 4 and st. 10 due to insufficient sample size) performed on the Bray-Curtis distances between samples confirmed that location has a significant effect on symbiont community (p = 0.001, R^2^ = 0.47 after 999 permutations). Host SV also had a significant effect (p = 0.009, R^2^ = 0.19), but only SV 5 was significantly different in pairwise comparisons. Analyses were performed with R packages phyloseq and vegan and plot was rendered with R package ggplot2.

### 3.2 Comparison of intra-host and free-living symbiont communities

To determine whether relative abundance of symbionts within hosts is a reflection of relative abundance of available symbionts in the surrounding water, we compared symbiont communities within acantharians collected from cruise stations in the ECS to environmental samples taken at the same time and place. 1,852,276 sequences were generated from environmental samples with 93,291– 246,564 reads per sample. After quality filtering and feature table construction, 645,325 sequences remained, with 35,691–89,163 sequences per sample. A PCoA plot based on Bray-Curtis distances between the entire community for each environmental sample confirmed that replicates from each location were more similar to each other than to replicates from other locations (Figure 6). Symbiotic SVs identified in acantharian samples comprised 1–3% of sequences in environmental samples and only 11 of 21 symbiotic SVs were also found in environmental samples (SVs 2, 5, 7, 11, 13, 14, 15, 16, 18, 19 were missing) (Figure 7). Intra-host and environmental symbiont communities clustered separately in a PCoA based on Bray-Curtis distances between samples (Figure 8) and the observed difference between community compositions in the two sample types was statistically significant (p = 0.001, R^2^ = 0.33). These results indicate that the intra-host community composition of symbiotic SVs is distinct from the surrounding environmental community composition.

**Figure 6.**
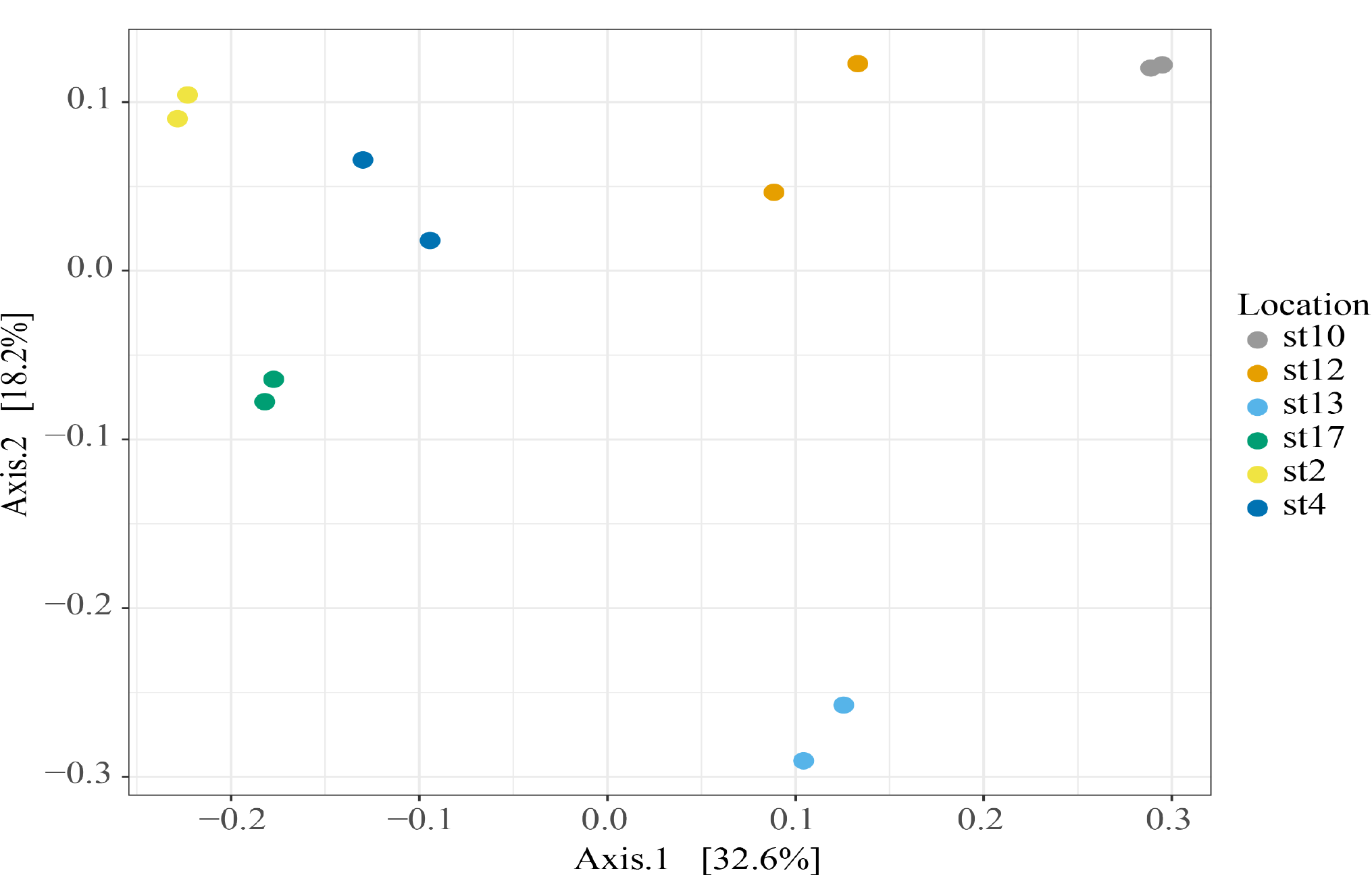
Biological replicates of environmental microbial eukaryote communities in each location are more similar to each other than replicates from other locations. PCoA plot from Bray-Curtis distances between ECS cruise station samples calculated from entire microbial eukaryote communities (> 0.2 μm, < 10.0 μm) in environmental samples collected from ECS cruise stations in May and June 2017. PCoA was performed with the R package phyloseq and plotted with ggplot2.

**Figure 7.**
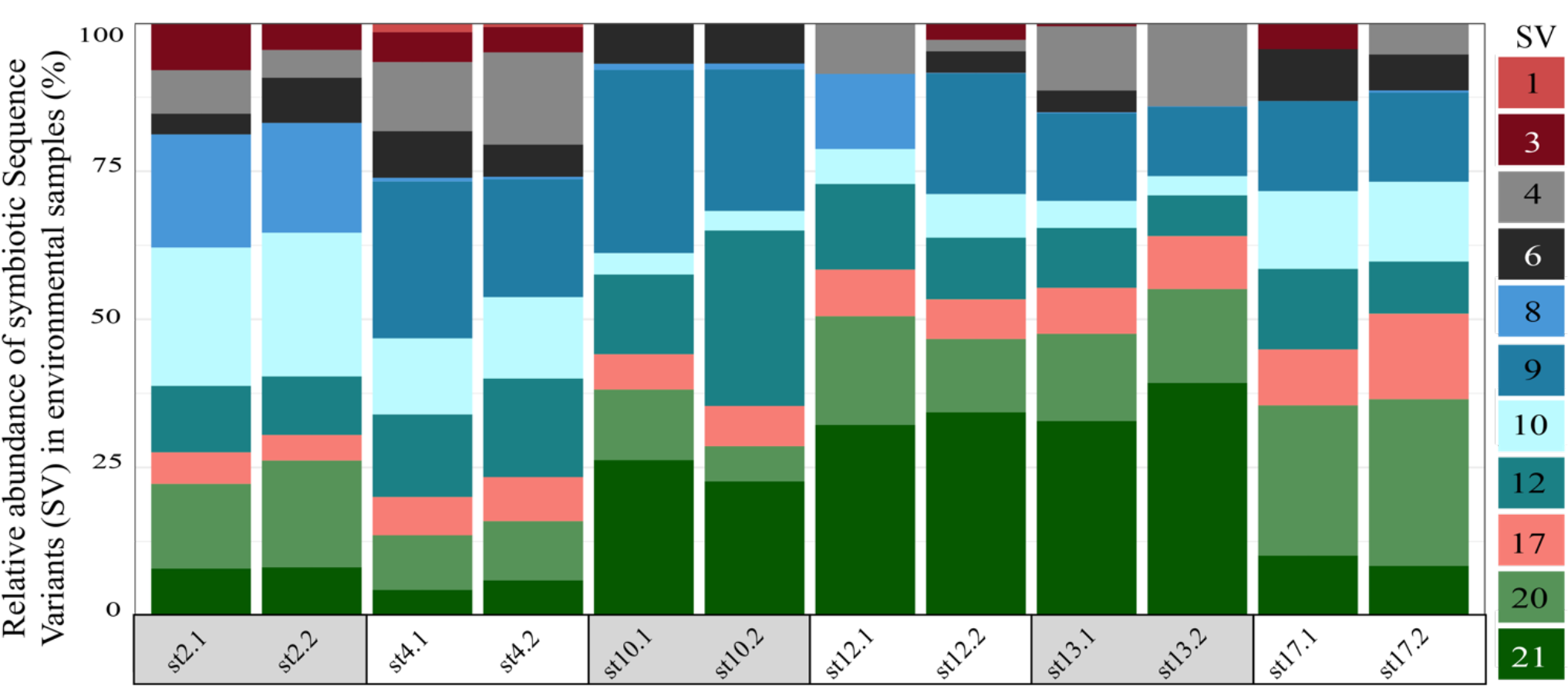
Relative abundances of symbiotic Haptophyte SVs in East China Sea (ECS) environmental samples are different than in ECS acantharian samples. The stacked bar graph displays the relative abundances of symbiotic SVs identified from acantharian samples that were also found in replicate environmental samples from each ECS cruise station. Symbiotic SVs are colored by clade: green is *Chrysochromulina*, blue is *Phaeocystis* clade Phaeo2, pink is *Phaeocystis cordata*, grey/black is *Phaeocystis jahnii*, and purple is *Phaeocystis*, but not placed with known clades. Only 11 of the symbiotic SVs were also found in environmental samples. *Chrysochromulina* SVs are present at all ECS cruise stations despite only being found in 1 acantharian sample from an ECS cruise station. The figure was created with R package ggplot2.

**Figure 8.**
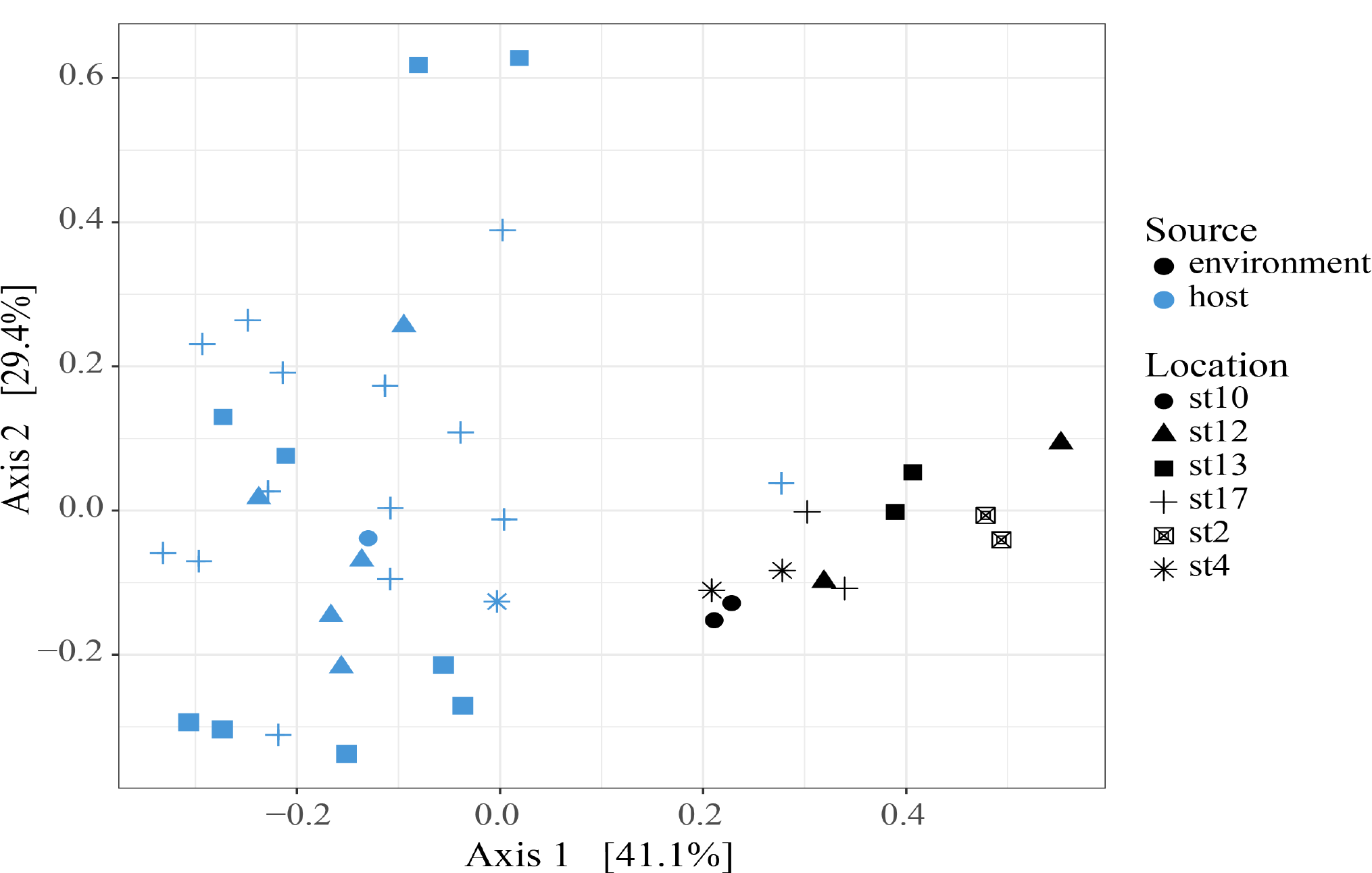
East China Sea (ECS) intra-host and free-living symbiont communities form distinct clusters in a Principal Coordinate Analysis (PCoA) plot. PCoA plot from Bray-Curtis distances between ECS environmental free-living and the host-associated symbiotic community compositions, including all symbiotic SVs identified in this study. The two sample types cluster separately and the difference in community composition is statistically significant (PERMANOVA, p = 0.001, R^2^ = 0.326, 999 permutations). Analyses were performed with R packages phyloseq and vegan and plot was rendered with R package ggplot2.

### 3.3 Visualization of host-associated symbionts and host digestive-organelles

By imaging symbiont chlorophyll autofluorescence with laser confocal microscopy, we were able to clearly enumerate photosynthetic symbionts within hosts. This technique allowed us to discern that there were more individual symbiont cells than observed SVs in each imaged acantharian collected near Okinawa in April and May (Figure 9A–B, Supplementary Figure S3). Symbionts with the free-living phenotype (twin parietal plastids, cell diameter < 5 μm) and symbionts with the symbiotic phenotype (more than 2 plastids, enlarged central vacuole, cell diameter > 5 μm) were observed inside the same host cells (Figure 9B, Supplementary Figure S3). By selectively staining host digestive-organelles with a fluorescent dye, we were able to visualize host lysosomes and phagolysosomes and their proximity to symbionts (Figure 9C–E). We observed lysosomes in both host exoplasm and endoplasm, while symbionts were only observed in host endoplasm. No symbionts were enclosed in phagolysosomes and lysosomes did not appear to be concentrated near symbiont cells.

**Figure 9.**
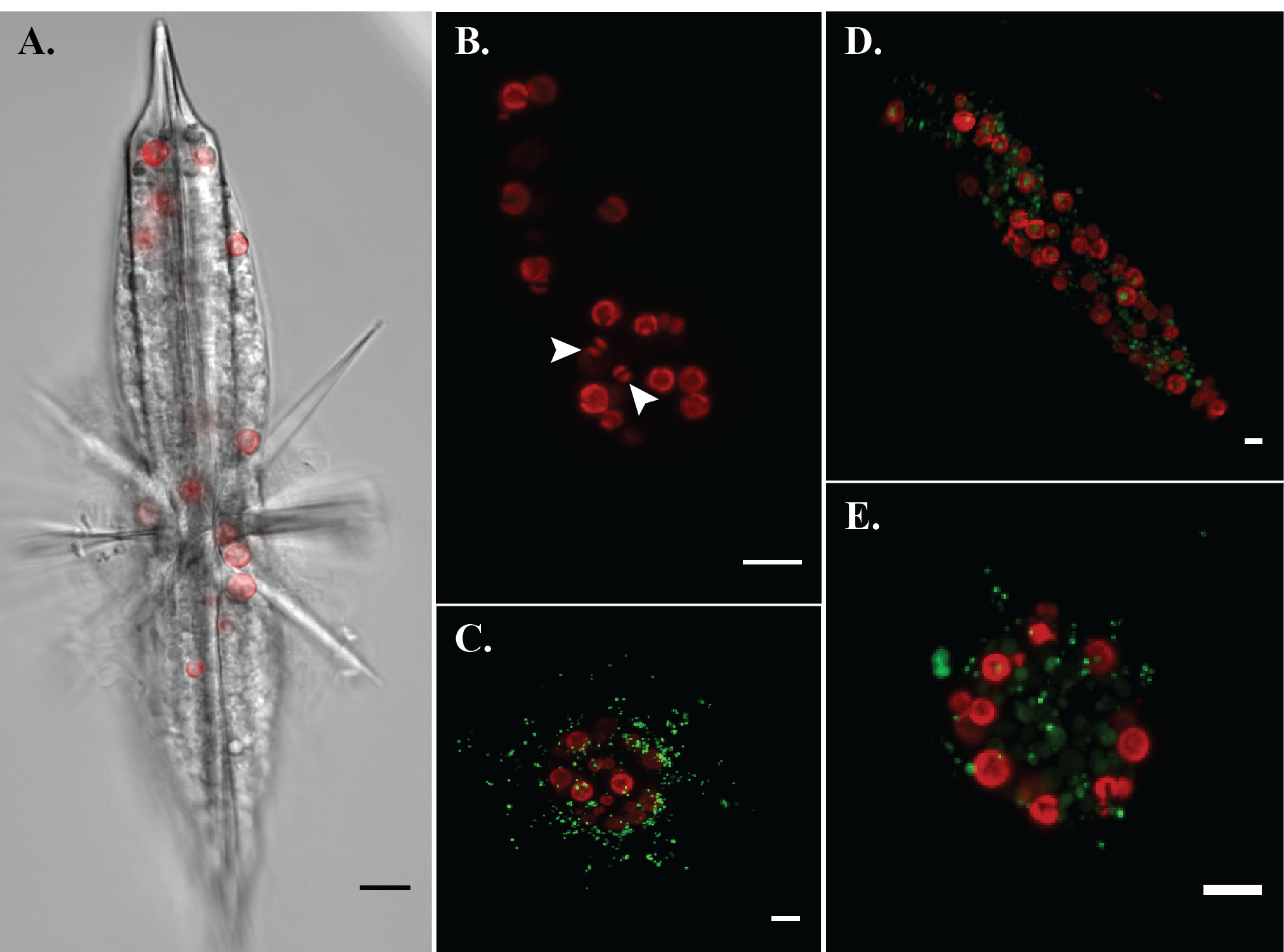
Fluorescent confocal microscopy of acantharians and their symbionts demonstrates more individual symbionts inside individual hosts than observed SVs and demonstrates that symbionts are not enclosed in digestive phagolysosomes. **A**. Single optical slice displaying autofluorescence of symbiont chlorophyll (red) and halogen light imaging of an entire host, sample Oki.3A. **B.** Maximum projection of a z-stack spanning the entire host sample Oki.4A, which contains 23 visible symbionts and 9 symbiotic sequence variants. Red fluorescence is symbiotic chlorophyll autofluorescence. Arrows indicate symbionts presenting the free-living phenotype: smaller cell diameter and two elongate, parietal chloroplasts. **C–E.** Maximum projections of z-stacks spanning entire hosts collected near Okinawa in December 2017. Red fluorescence is symbiotic chlorophyll autofluorescence. Green fluorescent staining is LysoTracker Green, which selectively binds to low-pH digestive organelles, including lysosomes and phagolysosomes. Symbionts are not held in phagolysosomes and lysosomes are not concentrated around symbionts, indicating that symbionts are not actively being digested. Scale bars are 10 μm in all panels.

## 4 Discussion

Acantharians are abundant globally and are especially abundant in low-nutrient subtropical gyres where they contribute to primary production by harboring intra-cellular algal symbionts (Michaels 1991; Michaels et al. 1995). Despite their ecological importance, host-symbiont dynamics in acantharian photosymbioses remain largely unstudied. Decelle et al. (2012) discovered that Acantharea-*Phaeocystis* symbioses are flexible in regard to symbiont species and hypothesized that the relationship is more akin to enslavement than mutualism (Decelle 2013; Decelle et al. 2012). If symbionts are exploited to the extent that they are unable to reproduce, hosts need to continuously recruit symbionts and should host diverse symbiont communities reflecting the relative availability of different symbionts. Our amplicon sequencing results show that acantharians do simultaneously host multiple species of *Phaeocystis* as well as *Chrysochromulina spp*., but we found that the host-associated symbiont community does not simply mirror the free-living community. Our microscopy results show that individual hosts harbor symbionts exhibiting the free-living phenotype as well as the symbiotic phenotype and demonstrate that symbionts are not being systematically digested. Together, these results suggest that hosts continue to recruit new symbionts, but they do not digest or shed symbionts at the same rate that they acquire them.

As acantharians increase in size, the number of symbionts they host also increases (Michaels 1991). To accomplish this, acantharians could recruit microalgal partners early in development and nurture reproducing symbiont populations. Alternatively, acantharians could recruit symbionts continuously, which would likely lead to multiple species of symbionts coexisting within individual hosts. In this study, all 42 acantharians collected hosted multiple species of *Phaeocystis* symbionts and several also hosted *Chrysochromulina spp*., suggesting that acantharians recruit symbionts more than once. The observation of symbionts with the free-living phenotype alongside symbionts with the symbiotic phenotype within single hosts is further evidence that hosts continue to recruit new symbionts (Febvre et al. 1979, Figure 9). However, these results do not exclude the possibility that symbionts reproduce *in hospite*, and symbionts are known to reproduce within larger photosymbiotic Rhizarians that can be maintained in culture (Takagi et al. 2016). Acantharians in this study contained more individual symbiont cells than unique Sequence Variants, indicating that either hosts recruit multiple symbiont cells from a clonal environmental population or that symbionts reproduce *in hospite*. Therefore, it is possible that symbiont cells exhibiting the free-living phenotype within hosts are recently divided cells rather than recently engulfed cells.

Since acantharians in this study simultaneously hosted multiple *Phaeocystis* and *Chrysochromulina* species, we expected that the intra-host symbiont community would reflect environmental symbiont availability. Our results, however, showed that the intra-host symbiont community was distinct from the free-living community. Additionally, acantharians sometimes hosted *Phaeocystis* genotypes (e.g. *P. jahnii* SV 5, *P. globosa* SVs 13 & 14) that were absent in environmental samples collected from the same place. These results suggest acantharians may maintain symbionts for extended periods, beyond that required for external populations to respond to changing environmental conditions. This is feasible since *Phaeocystis* generation times can be as brief as 6.6 h and vary by species in different light and temperature conditions (Jahnke 1989), while acantharians likely survive for at least a month (Suzuki & Not 2015). Intra-host communities may therefore be cumulative representations of all encountered environmental symbiont communities, rather than just a snapshot of the current community.

The observed differences between acantharians symbiont communities and the availability of symbionts could also indicate that hosts selectively uptake symbionts that are most beneficial in prevailing environmental conditions. *Chrysochromulina* SVs were dominant in each acantharian sample collected near Catalina Island, but were only found in one acantharian from an ECS cruise station, despite being well-represented in all cruise station environmental samples. Perhaps *Chrysochromulina spp*. make better partners in the Catalina Island environment compared to the Ryukyu Archipelago. If acantharians select and concentrate environmentally rare symbionts, it would also increase the likelihood that our environmental sampling missed those symbionts, especially because symbiotic SVs constituted only 1-3% of the community. Additionally, it remains possible that the differences observed between intra-host and environmental symbiont communities could derive from different nucleic acid extraction methods used. However, we did not observe digestion of symbionts by hosts, which provides additional support for the extended maintenance of symbionts *in hospite*.

This study demonstrates for the first time that intra-cellular symbiont diversity exists in cade F acantharian photosymbioses. Acantharians in the clade B genus*, Acanthochiasma*, also host multiple symbiont types, including *Chrysochromulina spp*. and several dinoflagellate genera (Decelle et al. 2012c). Radiolarian and foraminiferan (Rhizaria) hosts also host dinoflagellates and prasinophytes or prymnesiophytes (Gast & Caron 2001). Theoretically, simultaneously hosting multiple symbiont species or genotypes is ineffective and should negatively impact host fitness since different symbionts would compete for space and resources within hosts (Douglas 1998). Planktonic hosts can be transported long distances and may therefore experience larger environmental gradients on shorter time-scales than stationary photosymbiotic hosts. This could make hosting a diverse community of symbionts more effective for planktonic hosts, especially if some symbionts perform better under different conditions. Indeed, different *Phaeocystis* species have different light and temperature optima (Jahnke 1989) and different strains of a single species have varying abilities to utilize different nitrogen sources (Wang et al. 2011). Drifting buoys in the Global Drifter Program (Lumpkin et al. 2013) passing our sampling sites in spring, the season we sampled, traveled an average of 3.66° latitude in 30 days (n = 42), the estimated minimum survival time of acantharians (Suzuki & Not 2015), suggesting they can travel at least this far in their lifetime. While the associated mean temperature gradient of 1.93°C is smaller than experimental temperature gradients shown to differentially influence *Phaeocystis spp*., changes in day length and irradiance may affect photosynthetic output of *Phaeocystis* strains differently (Jahnke 1989).

The possible fitness benefit for symbionts associated with acantharians remains enigmatic, but the evidence we present for extended symbiont maintenance allows that *Phaeocystis* could glean some advantage from the symbiosis. Acantharian symbionts were not enclosed in phagolysosomes in host cells imaged in this study, which could be due to symbionts escaping phagosomes (Jamwal et al. 2016) or a failure of host lysosomes to fuse with symbiont-containing phagosomes (Hohman et al. 1982; Sibley et al. 1985). Electron microscopy suggests that there is a host membrane surrounding acantharian symbionts (Febvre et al. 1979), so it is more probable that lysosomes do not fuse with symbiont-containing phagosomes and they instead represent symbiosomes or long-term perialgal vacuoles. Symbiont signaling prevents lysosomes from fusing with symbiosomes housing *Chlorella* symbionts in *Paramecium* (Kodama et al. 2011) and *Hydra* (Hohman et al. 1982) hosts. Complex signaling between hosts and symbionts leading to development of symbiosomes and symbiont differentiation is suggestive of co-evolution (Hinde & Trautman 2001). If *Phaeocystis* also actively prevents lysosomes from fusing with phagosomes, then *Phaeocystis* has adapted to avoid digestion and perhaps to promote a stable symbiosis. Although our results suggest that acantharians maintain symbionts, we cannot rule out that they digest symbionts when stressed or before gametogenesis, which would rule out mutualism. Acantharian gametes do not contain symbionts, but it is not yet clear whether symbionts are digested or released during reproduction (Decelle & Not 2015). Additionally, it remains an open question whether released symbionts are reproductively competent outside the host (Decelle et al. 2012).

Our results demonstrate that clade F acantharians simultaneously associate with multiple *Phaeocystis* and *Chrysochromulina* species, providing further evidence that the Acantharea-*Phaeocystis* photosymbiosis is highly flexible. Our results suggest that symbionts escape host digestion for extended periods, but whether symbionts are capable of reproducing *in hospite* or after release, and whether they benefit from the relationship is still undetermined. Until acantharians can be maintained for prolonged periods under laboratory conditions, it will remain challenging to elucidate many aspects of the host-symbiont dynamics in this system. LysoTracker dyes can be utilized to track symbiosome conditions for as long as hosts survive and species- or genotype-specific fluorescent probes may be used to investigate whether different species or genotypes are differentially transformed into the symbiotic phenotype or are compartmentalized within the host. If symbionts with a single 18S rRNA gene Sequence Variant (or a unique combination of SVs) are co-localized, it would provide evidence for *in hospite* reproduction. Efforts to culture symbionts from *Phaeocystis*- hosting acantharians should continue. If successful, it would demonstrate unequivocally that symbionts maintain reproductive capacity and cultured symbionts would be an invaluable resource for comparative genomics and transcriptomics. Differential gene expression analysis could then be utilized to investigate physiological shifts in symbiotic *Phaeocystis* or *Chrysochromulina* compared to cultured free-living cells and could illuminate mechanisms of host-symbiont interaction. Further investigation into whether symbionts benefit from this relationship will be important to understanding host-symbiont dynamics in this and other protistan photosymbioses.

## 5 Author contribution

MB, MG, and SM planned the research. MB performed the research. LM optimized single-cell RNA techniques and performed research. MB analyzed the data and wrote the manuscript. MG and SM edited drafts.

## 6 Funding

This work was supported by funding from the Marine Biophysics Unit of the Okinawa Institute of Science and Technology Graduate University. MB is supported by a JSPS DC1 graduate student fellowship.

## 7 Conflict of Interest

The authors declare that the research was conducted in the absence of any commercial or financial relationships that could be construed as a potential conflict of interest.

## 8 Acknowledgements

We thank the captain and crew of the JAMSTEC R/V *Mirai* for their assistance and support in sample collection. Hiromi Watanabe, Dhugal Lindsay, and Yuko Hasagawa were instrumental in organizing and facilitating cruise sampling. Otis Brunner assisted in filtering seawater samples. We are grateful to David Caron for helpful discussion and instruction and for initiating and organizing sampling from Catalina Island. Alyssa Gellene further assisted in collecting and processing Catalina Island samples. Jo Tan assisted in sequencing library preparation and Hiroki Goto and the OIST sequencing section provided guidance and performed sequencing. Bassem Allam and Emmanuelle Pales-Espinosa gave invaluable feedback and suggestions. Members of the OIST Marine Biophysics Unit, Steven D. Aird, and David Caron edited and provided comments on the manuscript.

## 9 Data availability statement

Sequences generated and analyzed in this study have been submitted to the European Nucleotide Archive under the study accession number PRJEB24538. Intermediate data files and data analysis pipelines are available at https://github.com/maggimars/AcanthareaPhotosymbiosis and https://maggimars.github.io/AcanthareaPhotosymbiosis/Analysis.html.

